# Polarization effects and the calibration of a donut beam axial optical tweezers for single-molecule force spectroscopy

**DOI:** 10.1101/147967

**Authors:** Russell Pollari, Joshua Milstein

## Abstract

Advances in light shaping techniques are leading to new tools for optical trapping and micromanipulation. For example, optical tweezers made from Laguerre-Gaussian or donut beams display an increased axial trap strength and can impart angular momentum to rotate a specimen. However, their application to precision, biophysical measurements remains limited as their are a number of challenges to applying this tool to optical force spectroscopy. One notable complication, not present when trapping with a Gaussian beam, is that the polarization of the trap light can significantly affect the tweezers’ strength as well as the precise location of the trap. In this article, we provide a practical implementation of a donut beam optical tweezers for applying axial forces. We show how to precisely calibrate the height of the optical trap above the coverslip surface while accounting for focal shifts in the trap position that arise due to radiation pressure, mismatches in the index of refraction, and polarization induced intensity variations.

## INTRODUCTION

Optical tweezers are a versatile instrument for applying forces to the microscopic world and have emerged as the most precise tool for performing force spectroscopy experiments on biological molecules [1, 2]. Most optical tweezers make use of a tightly focused Gaussian beam; however, advancing methods in focal spot engineering have led to a range of novel optical traps, heralding a new generation of optical tweezers [3]. For instance, donut beams (also known as vortex beams or Laguerre-Gaussian beams) can be used to apply optical torques [4], and improve the axial trapping efficiency by reducing the radiation pressure along the optical axis [5–7]. While these unique optical traps can be directly applied to micromanipulation, there remain challenges to translating them into useful tools for precision force spectroscopy. Here we focus on developing donut beam optical tweezers into a precision single-molecule tool for performing axial force spectroscopy (i.e., along the direction of laser propagation) [8] and propose a number of innovative applications for such a device.

A Gaussian laser can be readily converted to a “donut-like” beam by introducing a spiral phase plate into the beam path of the form:

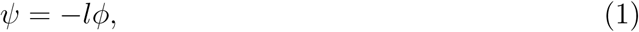

where *ϕ* is the azimuthal angle and l is an integer representing the topological charge. The result is a helical wave front with a phase singularity at the centre that produces a dark region along the optical axis. A more flexible way to introduce a spiral phase is with a phase-only spatial light modulator (SLM); the advantage being that the SLM can dynamically adjust the imprinted phase.

However, when focusing such a beam through a high-numerical aperture (N.A.) objective, polarization effects can significantly affect the intensity and phase profile [9, 10]. Numerical simulations of the tight focusing of a donut beam though a high-N.A. objective, using vectorial Debye theory, reveal that the presence of the characteristic dark central core depends on the polarization of the beam [11–13]. When the polarization of the beam has the same handedness as the topological charge of the phase, the intensity in the centre of the beam goes to zero. However, for linearly polarized light, the central dark spot begins to fill with light, and when the phase and polarization are anti-aligned, the intensity in the centre fills in significantly.

While these polarization effects are not visible in the far-field, so are difficult to directly image, we demonstrate their influence in altering the axial trapping strength. Jeffries et al. [13] previously showed that the axial trap strength of a donut beam tweezers is affected by the choice of right- or left-handed circularly polarized light. However, the axial trap strength is also dependent upon the location of the trap above the coverslip, and changing the polarization of the trap laser will alter the laser pressure, shifting the axial trap position. In this manuscript we explain how to correct for these effects. We show how surface effects can be used to precisely locate the height of the optical trap as the trap is moved axially above the plane of the coverslip, while accounting for the focal shift (in trap position) that arises due to radiation pressure, mismatches in the index of refraction, and polarization induced intensity variations. As expected, trapping with circularly polarized light aligned with the topological charge showed an increase in axial trapping efficiency compared to light of the opposite chirality. Our method of calibrating the precise strength and position of the optical trap lays the foundation for a new approach to axial force spectroscopy [8] via a donut beam optical tweezers.

## MATERIALS AND METHODS

### A. Optical Setup

The optical setup is shown in Fig 1. A 1062nm Nd:YAG laser (4W, Coherent BL-106C) incident on a phase-only spatial light modulator (SLM) (Hammamatsu X10468) is 4-f imaged onto the back focal plane (BFP) of an oil-immersion objective (Olympus PlanApo 100x, 1.4 NA). The laser is vertically polarized using a half wave plate (HWP) and a polarizing beam splitter (PBS) which also allows for manual tuning of the laser power. An iris placed in an intermediate Fourier plane blocks the zeroth order (unmodulated) and any unwanted higher-orders of light reflected off the SLM. The light can be circularly polarized by inserting a quarter wave plate (QWP) before focusing into the sample. A condensor (Olympus LUCPLanFL 40x, 0.75 NA) collects the scattered light and images it onto a position sensitive diode (PSD) (FirstSensor DL100-7-PCBA3) for back focal plane interferometry (BFPI). The axial position of the laser focus z is controlled by superimposing a Fresnel lens to the hologram:

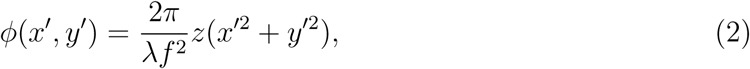

where *x*′, *y*′ are the pixel coordinates on the SLM, *λ* is the laser wavelength, and *f* is the focal length of the objective.

**FIG. 1.**
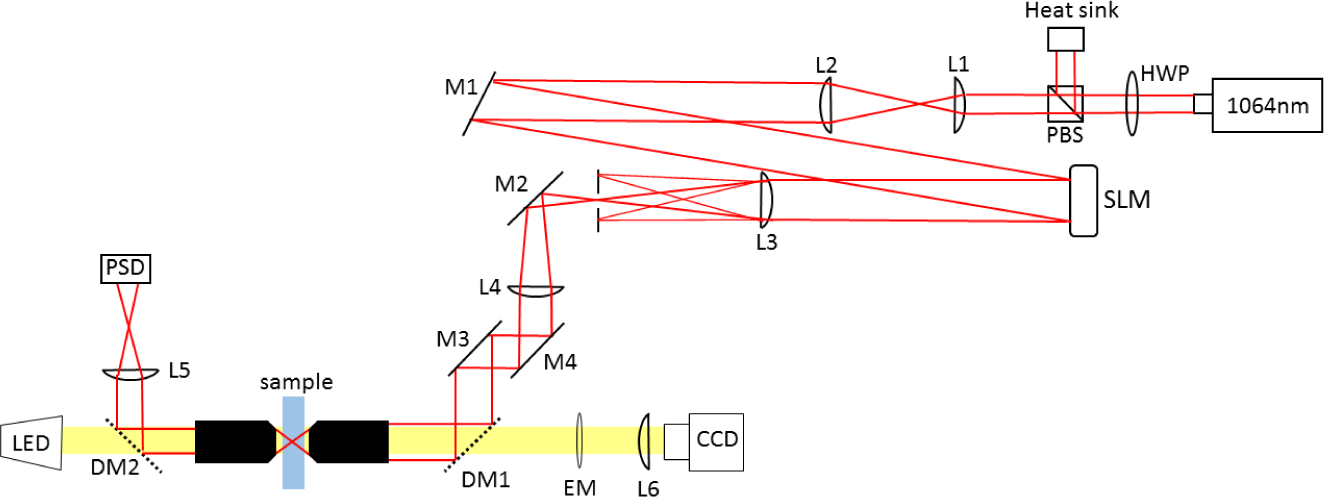
Optical Setup. A 1064 nm, near-infrared laser is projected onto a phase-only SLM, and imaged on the back aperture of a high-N.A. objective. An aperture is introduced shortly after the SLM to block all but the first-order diffracted light. The sample plane is illuminated by a white-light LED and imaged both via a CCD and through back focal plane interferometry onto a position sensitive photodiode (PSD).

To create a donut beam, Eq. (1) is superimposed on the SLM with a blazed grating designed to displace the beam a distance x from the optical axis, ensuring no interference from unmodulated light reflected off the SLM. The superposition is often referred to as a forked grating:

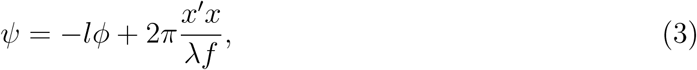

where *x*′ is the SLM coordinate, *λ* is the wavelength, and *f* is the focal length of the Fourier transforming lens.

Introducing the phase above generates a field distribution in the Fourier plane that is actually a superposition of radial, higher-order Laguerre-Gaussian modes 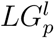, which results in multiple rings around a central dark spot (see Appendix A). There are various approaches to rectifying this issue. For instance, by spatially tuning the efficiency of a blazed phase grating, a phase-only SLM can effectively encode amplitude information [14]. This additional control can be used to modulate all but the lowest order 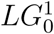 mode to obtain an essentially pure donut beam; however, we found that purifying the donut beam resulted in a significant reduction in the trap strength (~ 50% or more) due to a loss of intensity. For our measurements, we chose to neglect the mode purity corrections to maintain a sufficient trap strength. Since, absent these corrections, the light is already primarily in the 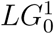 mode, this approximation did not qualitatively affect our results.

### B. Polarization Effects

When tightly focusing light with a spatially varying phase profile, such as a donut beam, the polarization of the light can strongly impact its intensity profile at the focus. To better understand the effects of polarization on a donut beam optical trap, the tight focusing of a donut beam though a high-N.A. objective was investigated using vectorial Debye diffraction theory [15]. This approach properly treats the polarization of the trap light, which is not accounted for in scalar diffraction theory (see Appendix B).

Figure 2 shows the axial intensity profile, numerically computed in MATLAB, of the focused laser light for these three polarizations (RHC, LHC, and Linear) as well as intensity cross sections through the centre of each optical trap. When the polarization has the same handedness as the topological charge of an 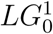 beam, the central intensity truly goes to zero. Here we have oriented the topological charge so that it is aligned with right-handed circularly (RHC) polarized light. However, when the handedness is of the opposite sign (LHC), the intensity in the centre fills in significantly. And a linearly polarized LG beam has a central intensity somewhere between these two extremes. This same effect is not prominent with lower N.A. objectives, but becomes increasingly important with tighter focusing as is necessary for optical trapping. Figure 3 shows the increasing need to account for polarization effects as the N.A. is increased. In the figure, we present the effects of RHC, LHC, and Linear orientations for increasing N.A. (N.A. = 0.1, 1.0 and 1.4). While at low N.A., the intensity profiles are independent of the polarization, as the N.A. increases, the intensity profiles become strongly dependent on the polarization. We note that these problems could be circumvented by employing more complex polarization schemes, such as azimuthally polarized light, which has been shown to maintain a dark core under tight focusing conditions [16]. The cost, however, is that one needs to generate this novel polarization state.

**FIG. 2.**
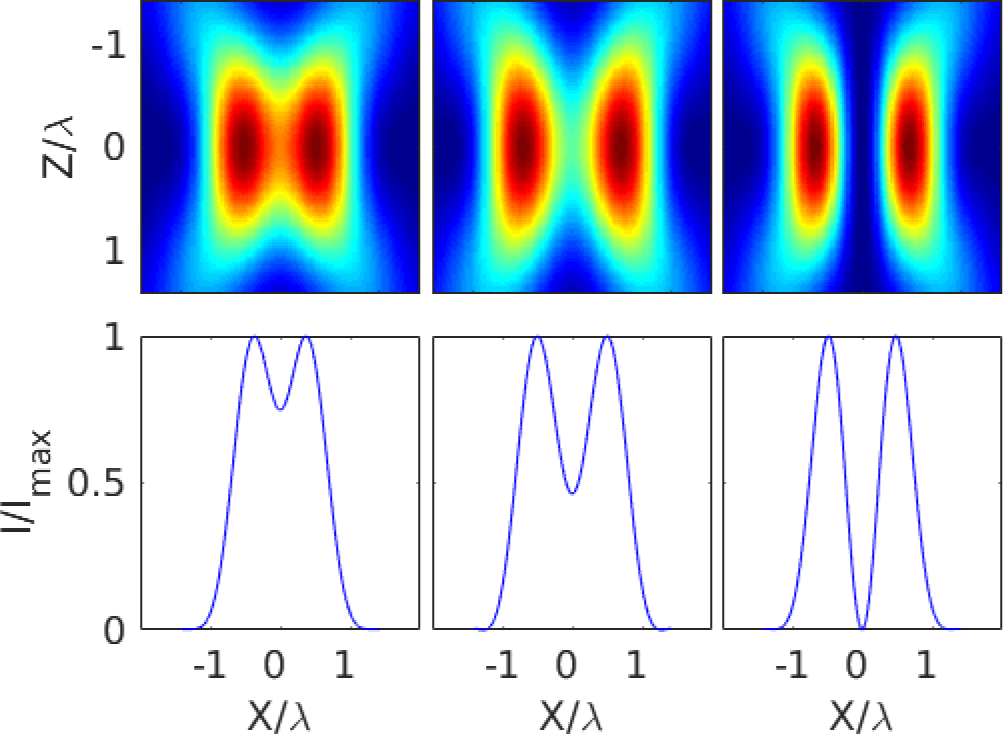
Polarization Dependent Focal Distribution. (Top) Cross-section of the focused light intensity profile along the axial laser direction (N.A. = 1.4). (Bottom) Normalized intensity through the centre of the focal spot. From left to right: LHC (anti-aligned), Linear, and RHC (aligned) polarized incident light.

**FIG. 3.**
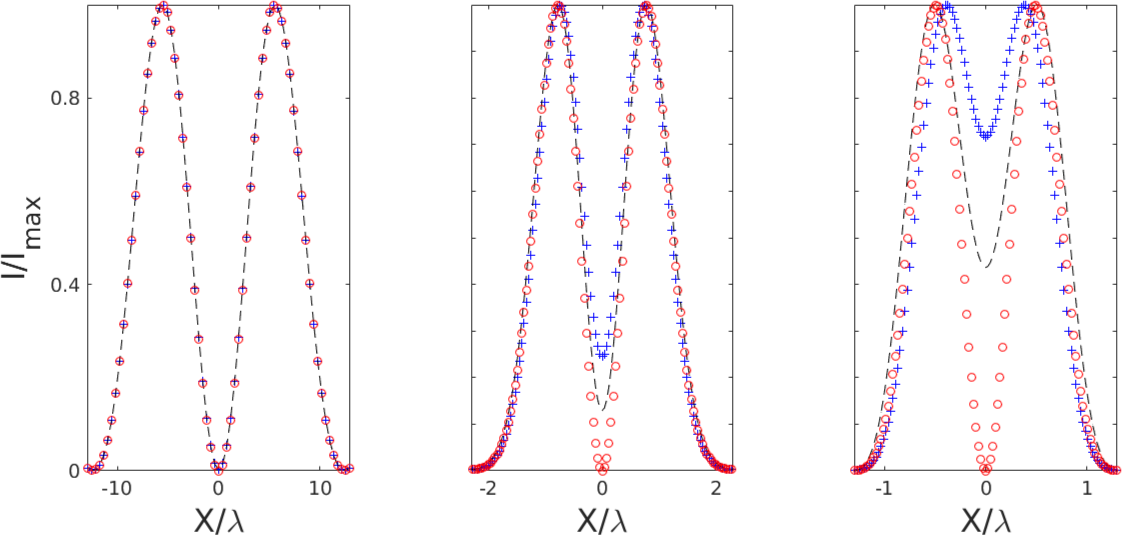
Polarization Dependence at Varying Focal Strength. Intensity profiles of 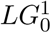 beams with (from left to right) N.A. = 0.1,1.0, and 1.4. Linear (dashed line), RHC (aligned, circles), and LHC (anti-aligned, crosses). As the N.A. increases, the intensity profile becomes increasingly dependent on the polarization.

### C. Relations for Optical Trap Calibration

The standard approach to measuring position within an optical tweezers is through backfocal-plane-interferometry (BFPI) [17]. The Gouy phase shift between light scattered by a trapped micro-bead and unscattered light results in an intensity variation at the back focal plane of the condensor. This variation is proportional to the trapped bead’s axial displacement relative from the trap centre *δz*(*t*), and when imaged on a photodiode is translated into a voltage signal *V*(*t*).

A measure of the detector sensitivity *β* can be used to convert from a voltage to a distance *δz*(*t*) = *βV*(*t*). Both the trap stiffness *κ* and the sensitivity *β* can be determined by measuring the power spectrum of an untethered, trapped bead over the relevant frequencies *f*. The power spectrum takes the form of a Lorentzian as follows [18]:

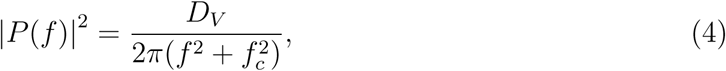

with the diffusivity *D_V_* (measured in volts) and the corner frequency *f_c_* extracted by fitting to the data. The trap stiffness and detector sensitivity are determined, respectively, by the relations:

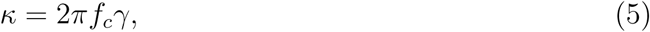

and
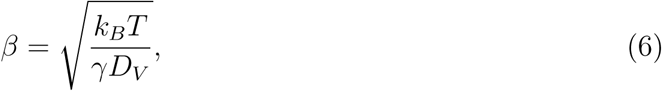

where *k_B_T* = 4.1 pN • nm and γ is the hydrodynamic drag coefficient. For a microsphere of radius *R*, the axial drag on the particle, close to the coverslip surface, can be approximated by Brenner’s formula:

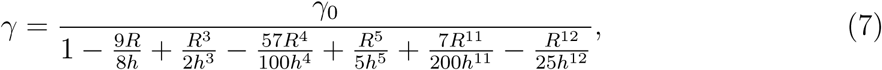

where *h* is the trap height, defined as the distance between the trapped bead’s centre and the coverslip, and *γ*_0_ = 6*πηR* is the drag in an infinite medium of viscosity *η* [19]. Equation 7 is clearly nonlinear, so when trapping close to the coverslip surface (roughly, *h* ≤ 3*R*), this calibration becomes acutely sensitive to the height of the trap. We will take advantage of this sensitivity later in calibrating the focal shift of the trap.

## RESULTS

Calibrating a donut beam optical tweezers for use in precision, axial force spectroscopy is challenging. First, due to the radiation pressure, the trap focus is shifted downstream of the laser focus. Second, in large part due to the mismatch in the indices of refraction between the oil/coverslip and aqueous trapping medium, the strength of the optical trap varies as a function of height, which is clearly problematic. We have previously shown that these challenges can be overcome for an axial optical tweezers generated from a Gaussian beam [20]. One can accurately measure the height of the trap from oscillations in the intensity signal due to multiply-reflected light (between a trapped microsphere and the coverslip surface) and even correct for the index mismatch by superimposing an additional hologram.

Unfortunately, a majority of the light that gives rise to the intensity oscillations is lost when working with donut beams, so much so that we are unable to simply apply our previous results of Ref. [20] to the present case. Fortunately, an alternate approach that makes use of surface interactions can be adapted to the current situation. That is, if we can correct for the index mismatch, so that the trap strength remains constant at varying depths above the coverslip, we can apply Brenner’s relation (Eq. 7) to accurately calibrate the height of the various donut traps.

In Ref. [20] we showed that correcting for first order spherical aberrations alone was sufficient to correct the index mismatch and achieve a constant axial trap stiffness. Since the correction is independent of the intensity profile, it should also be applicable for optical traps generated by donut beams. At each focal depth *z*, this correction can be imposed by displaying the following phase pattern on the SLM:

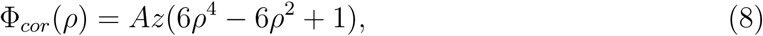

where *ρ* is the radial coordinate normalized to the objective’s entrance pupil radius and the constant *A* is empirically determined.

We initially followed this approach to achieve a constant axial trap stiffness for a Gaussian beam (with an *R* = 500 nm polystyrene microsphere), then converted to a circularly polarized donut beam by applying a forked grating to the SLM and inserting a *λ*/4 wave-plate before the objective. Figure 4 shows an experimental measure of the trap strength as a function of axial position for both RHC (aligned) and LHC (anti-aligned) light. With the phase correction of Eq. 8 applied, the trap stiffness for both orientations of the donut beam remain constant for at least 3 *μ*m above the coverslip surface. In fact, due to a reduced laser pressure, the axial trapping strength of the RHC polarized trap is ~ 44% stronger than that of the LHC trap. Since the trap stiffness *κ* remains constant as a function of the height *h* of the trap, *h* can be directly extracted from a measurement of the corner frequency at each axial position (Eqs. 5 and 7), which is how we obtained the horizontal axis in Fig 4. Relative spatial deviations *δz*(*t*) from this height can then be measured through standard BFP interferometry (Sec. C) to precisely track the axial location of a microsphere within a donut beam optical trap.

**FIG. 4.**
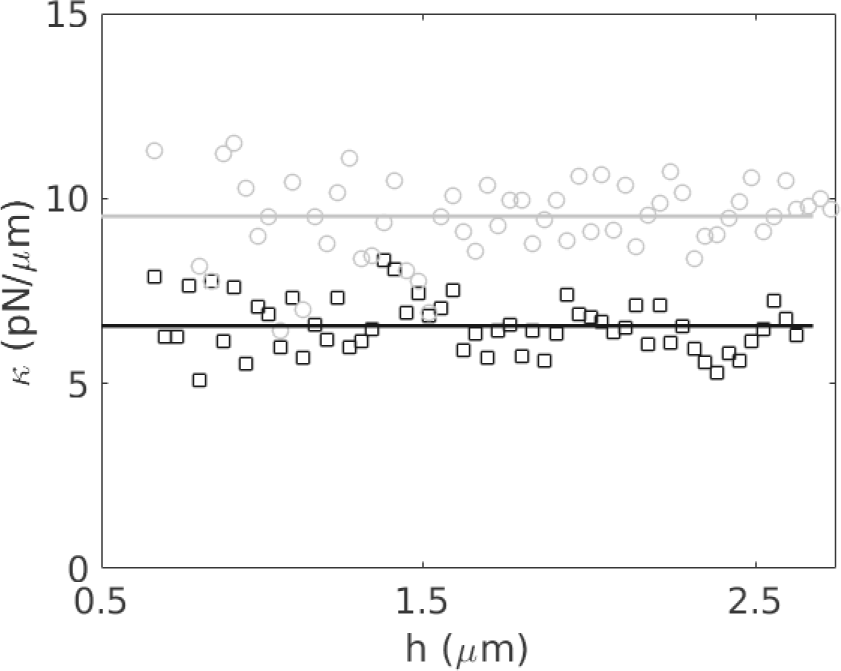
Polarization Dependent Trap Strength. Trap strength κ as a function of height *h*, displaying a constant trap stiffness, for LHC (anti-aligned, squares) and RHC (aligned, circles) polarized light. The solid lines are averages of the data at each height. The axial trap strength of the RHC (aligned) polarized trap is 1.44× stronger than the LHC (anti-aligned) polarized trap.

## DISCUSSION

We have shown that, when generating an optical tweezers with a donut beam, the trapping laser’s polarization must be considered with respect to the imprinted topological charge. We have found that the trap strength increases by a factor of 1.44× as the laser polarization goes from being anti-aligned to aligned with the topological charge. This can easily be achieved by simply rotating a quarter-wave plate (as we have done here) or, more rapidly, by employing a liquid crystal retarder. Such an approach would provide a way to apply subtle axial forces without a need to move either the stage or the laser focus, giving rise to a unique, new type of axial optical tweezers.

Another application of our results may be to aid in combining single-molecule fluorescence with optical tweezers. Single-molecule fluorescence can yield direct information on chemical kinetics and local structural changes. Using both techniques in tandem can provide significant new insight into molecular mechanisms inaccessible to either technique alone [21, 22]. Combining the two techniques is not trivial, however. The high intensity light from the trapping laser tends to increase the bleaching rate of fluorescent labels as well as obscure the significantly weaker fluorescence signal [23]. These issues are only significant near the focus of the trapping laser, so can be avoided with sufficient spatial separation of the trap from the fluorescent labels. One may also temporally separate the trapping and excitation beams as the nonlinear processes that drive the enhanced photobleaching are greatly reduced when the fluorophores are not excited [24]. Our work suggests a new approach, similar to the idea of employing donut beams to reduce photodamage proposed in [13], and that is simply to locate a fluorescently labeled molecule within the intensity minimum of the donut beam (with the laser polarization aligned to ensure a true, dark central region). Application of axial forces would maintain the label’s position within the centre of the donut beam. This would ensure that both the trap laser and fluorescence excitation laser are not incident on the labels simultaneously and should mitigate the issues with enhanced photobleaching.

## ACKNOWLEDGMENTS

R. Pollari and J. N. Milstein acknowledge support from the Natural Sciences and Engineering Research Council of Canada (NSERC, RGPIN 418251-13) and the Canada Foundation for Innovation (CFI, PN 30735).

## Appendix A Optimizing the Mode Purity

**FIG. 5.**
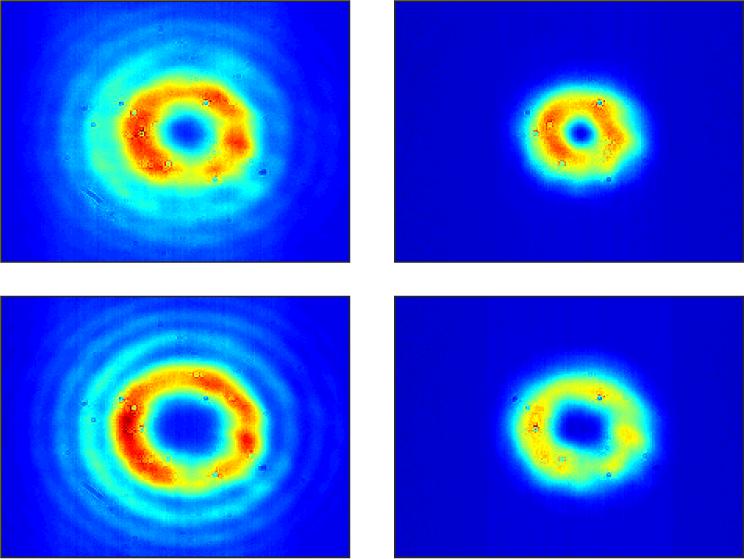
Far-field intensity profiles (Left) before and (Right) after modifying the hologram to encode amplitude information to purify the 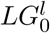 mode. The profiles for (Top) *l* = 1 and (Bottom) *l* = 2 are shown.

A Gaussian beam incident on a forked grating with topological charge *l* produces a field *U_l_* in the Fourier plane that is actually a superposition of Laguerre-Gaussian modes with azimuthal order *l* and radial orders *p*:

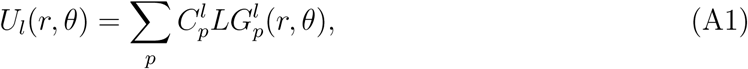

where 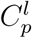 is a coefficient determining the amplitude of each mode and
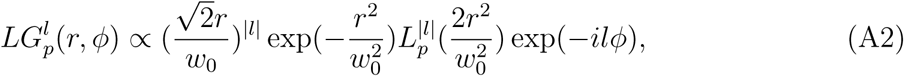

where *w*_0_ is the beam waist and 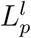 are generalized Laguerre polynomials [25]. For low values of *l*, the majority of the power ends up in the *p* = 0 mode, which corresponds to a single ring, but a portion of the intensity goes into higher radial modes [26, 27]. One approach to purifying the lowest mode is to encode amplitude information in the phase-only hologram. This can be achieved by spatially modulating the diffraction efficiency of the blazed grating imprinted on the SLM. There are a number of ways to implement this approach [27, 28]. One of the most efficient ways to encode the amplitude profile A, so as to maintain the intensity of the final hologram, is as follows:

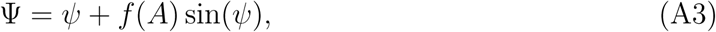

where *J*_0_ [*f*(*A*)] = *A* and *J*_0_ is the zeroth order Bessel function of the first kind [28]. Thus, single ring intensity profiles are produced by setting 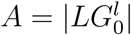. Figure 5 shows the intensity profiles for *l* = 1 and *l* = 2, before and after the addition of the amplitude mask. The higher order radial modes, seen as additional outer rings, vanish with the mask applied.

## Appendix B Vectorial Debye Theory

The electric field vector near the focus is given by:

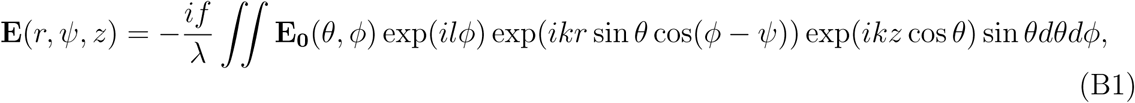

where *ϕ* is the azimuthal angle, and *θ* is the angle of the numerical aperture (see Fig 6). **E**_0_(*θ*, *ϕ*) is the electric field vector along a spherical surface S centred on the focal spot (see Fig 6). For circularly polarized light focused through an objective obeying the sine condition [15]:

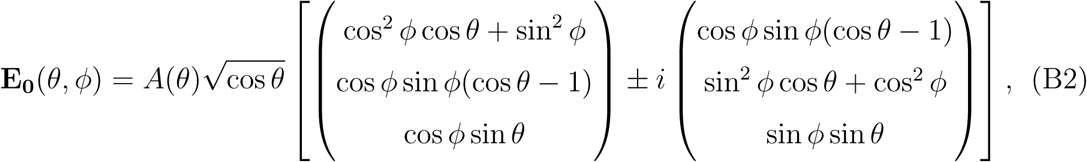

where ± denotes right-handed or left-handed circular polarization and *A*(*θ*) is the amplitude distribution along the surface *S*. Inserting Eq. B2 into Eq. B1 yields the electrical field component in all three dimensions:

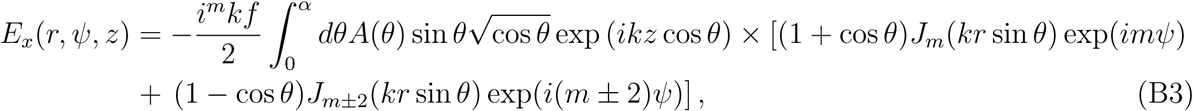

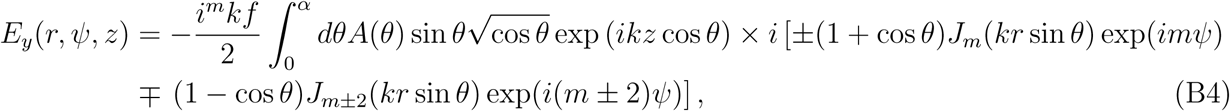

and
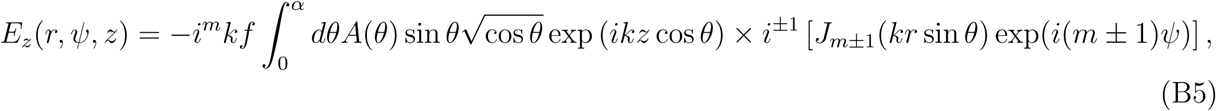

where *α* = sin^-1^(*N*.*A*./*n*) is the maximal angle of the objective imaged through a medium of refractive index *n*, and *J*_m_ are Bessel functions of the first kind.

**FIG. 6.**
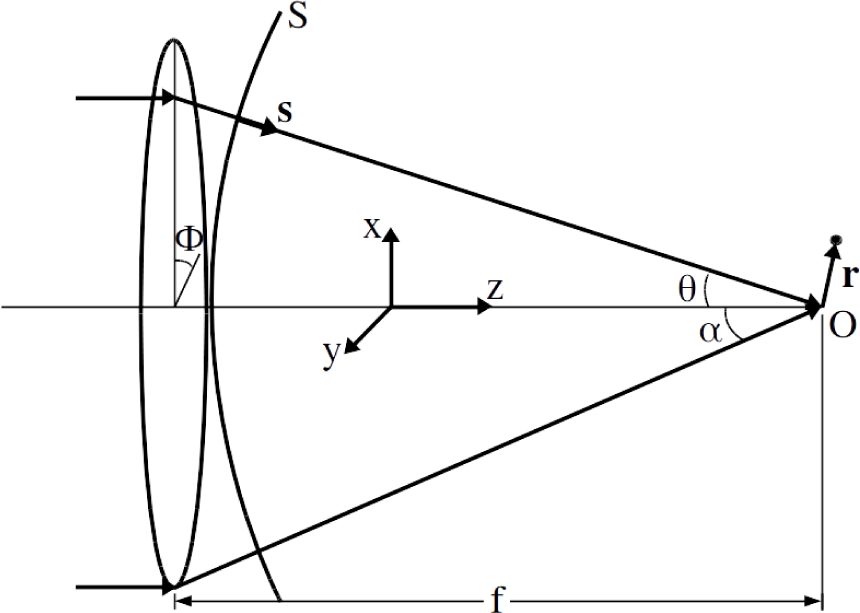
Geometry for numerical Debye simulations.

## References

[1] Moffitt JR, Chemla YR, Smith SB, Bustamante C. Recent advances in optical tweezers. Annu Rev Biochem. 2008;77:205–228.

[2] Heller I, Hoekstra TP, King GA, Peterman EJG, Wuite GJL. Optical Tweezers Analysis of DNA-Protein Complexes. Chem Rev. 2014;114(6):3087–3119.

[3] Dholakia K, Cizmar T. Shaping the future of manipulation. Nat Photonics. 2011;5(6):335–342.

[4] Simpson NB, Allen L, Padgett MJ. Optical tweezers and optical spanners with Laguerre-Gaussian modes. J Mod Opt. 1996;43(12):2485–2491.

[5] Friese MEJ, Rubinsztein-Dunlop H, Heckenberg NR, Dearden EW. Determination of the force constant of a single-beam gradient trap by measurement of backscattered light. Appl Optics. 1996;35(36):7112–7116.

[6] He H, Heckenberg NR, Rubinsztein-Dunlop H. Optical-particle trapping with higher-order doughnut beams produced using high-efficiency computer-generated holograms. J Mod Opt. 1995;42(1):217–223.

[7] Simpson NB, McGloin D, Dholakia K, Allen L, Padgett MJ. Optical tweezers with increased axial trapping efficiency. J Mod Opt. 1998;45(9):1943–1949.

[8] Yehoshua S, Pollari R, Milstein JN. Axial Optical Traps: A New Direction for Optical Tweezers. Biophys J. 2015;108(12):2759–2766.

[9] Zhao Y, Edgar JS, Jeffries GDM, McGloin D, Chiu DT. Spin-to-Orbital Angular Momentum Conversion in a Strongly Focused Optical Beam. Phys Rev Lett. 2007;99:073901.

[10] Chen B, Zhang Z, Pu J. Tight focusing of partially coherent and circularly polarized vortex beams. J Opt Soc Am A-Opt Image Sci Vis. 2009;26(4):862–869.

[11] Bokor N, Iketaki Y, Watanabe T, Fujii M. Investigation of polarization effects for highnumerical-aperture first-order Laguerre-Gaussian beams by 2D scanning with a single fluorescent microbead. Opt Express. 2005;13(26):10440–10447.

[12] Hao X, Kuang C, Wang T, Liu X. Effects of polarization on the de-excitation dark focal spot in STED microscopy. J Opt. 2010;12(11).

[13] Jeffries GDM, Edgar JS, Zhao Y, Shelby JP, Fong C, Chiu DT. Using polarization-shaped optical vortex traps for single-cell nanosurgery. Nano Lett. 2007;7(2):415–420.

[14] Davis JA, Cottrell DM, Campos J, Yzuel MJ, Moreno I. Encoding amplitude information onto phase-only filters. Appl Opt. 1999;38(23):5004–5013.

[15] Gu M. Advanced optical imaging theory. vol. v. 75. Berlin: Springer; 2000.

[16] Youngworth KS, Brown TG. Focusing of high numerical aperture cylindrical-vector beams. Opt Express. 2000;7(2):77–87.

[17] Gittes F, Schmidt CF. Interference model for back-focal-plane displacement detection in optical tweezers. Opt Lett. 1998;23(1):7–9.

[18] Berg-Sorensen K, Flyvbjerg H. Power spectrum analysis for optical tweezers. Rev Sci Instrum. 2004;75(3):594–612.

[19] Schaeffer E, Norrelykke SF, Howard J. Surface forces and drag coefficients of microspheres near a plane surface measured with optical tweezers. Langmuir. 2007;23(7):3654–3665.

[20] Pollari R, Milstein JN. Improved axial trapping with holographic optical tweezers. Opt Express. 2015;23(22):28857–28867.

[21] Hohng S, Zhou R, Nahas MK, Yu J, Schulten K, Lilley DMJ, et al. Fluorescence-force spectroscopy maps two-dimensional reaction landscape of the Holliday junction. Science. 2007;318(5848):279–283.

[22] Brau RR, Tarsa PB, Ferrer JM, Lee P, Lang MJ. Interlaced optical force-fluorescence measurements for single molecule biophysics. Biophys J. 2006;91(3):1069–1077.

[23] van Dijk MA, Kapitein LC, van Mameren J, Schmidt CF, Peterman EJG. Combining optical trapping and single-molecule fluorescence spectroscopy: Enhanced photobleaching of fluorophores. J Phys Chem B. 2004;108(20):6479–6484.

[24] Ferrer JM, Fangyuan D, Brau RR, Tarsa PB, Lang MJ. IOFF Generally Extends Fluorophore Longevity in the Presence of an Optical Trap. Curr Pharm Biotechnol. 2009;10(5):502–507.

[25] Karimi E, Zito G, Piccirillo B, Marrucci L, Santamato E. Hypergeometric-gaussian modes. Opt Lett. 2007;32(21):30533055.

[26] Sephton B, Dudley A, Forbes A. Revealing the radial modes in vortex beams. Appl Optics. 2016;55(28):78307835.

[27] Ando T, Ohtake Y, Matsumoto N, Inoue T, Fukuchi N. Mode purities of Laguerre-Gaussian beams generated via complex-amplitude modulation using phase-only spatial light modulators. Opt Lett. 2009;34(1):3436.

[28] Arrizon V, Ruiz U, Carrada R, Gonzalez LA. Pixelated phase computer holograms for the accurate encoding of scalar complex fields. J Opt Soc Am A 2007;24(11):3500 3507.

